# Use of the particle agglutination/particle agglutination-inhibition test for antigenic analysis of SARS-CoV-2

**DOI:** 10.1101/2022.09.08.507221

**Authors:** Jun Kobayashi, Shutoku Matsuyama, Masayuki Shirakura, Tomoko Arita, Yasushi Suzuki, Hideki Asanuma, Shinji Watanabe, Hideki Hasegawa, Kazuya Nakamura

## Abstract

The antigenicity of SARS-CoV-2 is a critical issue for the effectiveness of the vaccine, and thus it should be phenotypically evaluated by serological assays as new field isolates emerge. The hemagglutination/hemagglutination-inhibition (HA/HI) tests are well-known as a representative method for antigenic analysis of influenza viruses, but SARS-CoV-2 is unlikely to agglutinate to human or guinea pig red blood cells. Therefore, the antigenic analysis requires complicated enzyme-linked immunosorbent assay (ELISA) or cell-based assays such as the microneutralization assay. In this study, we developed the particle agglutination/particle agglutination-inhibition (PA/PAI) test to easily and rapidly quantify the virus and antibody using human angiotensin-converting enzyme 2 (hACE2)-bound latex beads. The PA titer was positively correlated with the plaque-forming units. The PAI titer using post-infection Syrian hamster antisera clearly revealed the antigenic difference between the omicron and previous variants. The results show the PAI test is useful for easy and rapid antigenic analysis of SARS-CoV-2.

## Introduction

SARS-CoV-2, which causes COVID-19, has infected over 500 million people and killed over 6 million, despite the production of over 11 billion doses of vaccine as of April 2022 (WHO COVID-19 Dashboard, https://covid19.who.int/). The pandemic of SARS-CoV-2, with its continuously evolving viral properties, has become a major public health concern that adds to the existing concern regarding influenza virus pandemics/epidemics. In particular, the omicron variants that emerged in November 2021 have a higher number of mutations than the previous variants and are considered to have relatively mild symptoms, but to be highly infectious. Furthermore, the omicron variants are resistant to immunity raised by previous variants^1,2^, and thus have caused the largest number of infections.

To predict the antigenicity and other viral properties of each isolate, a large number of the viral genomes have been sequenced using next-generation sequencers and registered (GISAID; https://www.gisaid.org/)^3^. However, the antigenicity of each variant must also be phenotypically evaluated by serological studies. The neutralization assay^4^ is one of the methods to analyze the antigenicity of SARS-CoV-2 isolates, but it is a time-consuming and complicated cell-based assay. A simpler enzyme-linked immunosorbent assay (ELISA) method without cells has also been reported^5^, but it requires artificially modified/purified proteins, such as the human angiotensin-converting enzyme 2 (hACE2) and the receptor binding domain of the SARS-CoV-2 spike protein, and a plate reader.

The antigenicity of influenza virus has been easily and rapidly analyzed by means of hemagglutination (HA)/hemagglutination-inhibition (HI) tests^6^. SARS-CoV-2 shows the hemadsorption activity with human erythrocytes on the virus-infected Vero cells, but does not exhibit direct HA activity using human and guinea pig erythrocytes^7^. In addition, there are increasing limitations on the use of blood cells due to various issues, such as animal ethics^8^. Therefore, the HA/HI tests are not applicable for SARS-CoV-2, and a surrogate for blood cells is required.

In this study, we established particle agglutination (PA)/particle agglutination-inhibition (PAI) methods using hACE2-bound latex beads as a surrogate for blood cells, utilizing the phenomenon that SARS-CoV-2 interacts with hACE2 *via* its spike protein^9^. These methods enable easy and rapid measurement of SARS-CoV-2 titer and antibody titer against the virus as well as antigenic analysis without special equipment.

## Results

## Establishment of a particle agglutination (PA) test

First, we aimed to establish a PA test as a virus titration method using hACE2-bound latex beads (hACE2-beads) as a surrogate for the blood cells used in the HA test of influenza viruses. The hACE2-beads were prepared based on the method used for SARS-CoV-2 antigen-coated latex beads^10^ (see the Online Methods). A suspension of the prepared beads showed a clear sedimentation pattern after overnight settling at a final concentration of 0.03% (Fig. 1a), which was used as the initial condition of the preliminary PA test.

**Figure 1.**
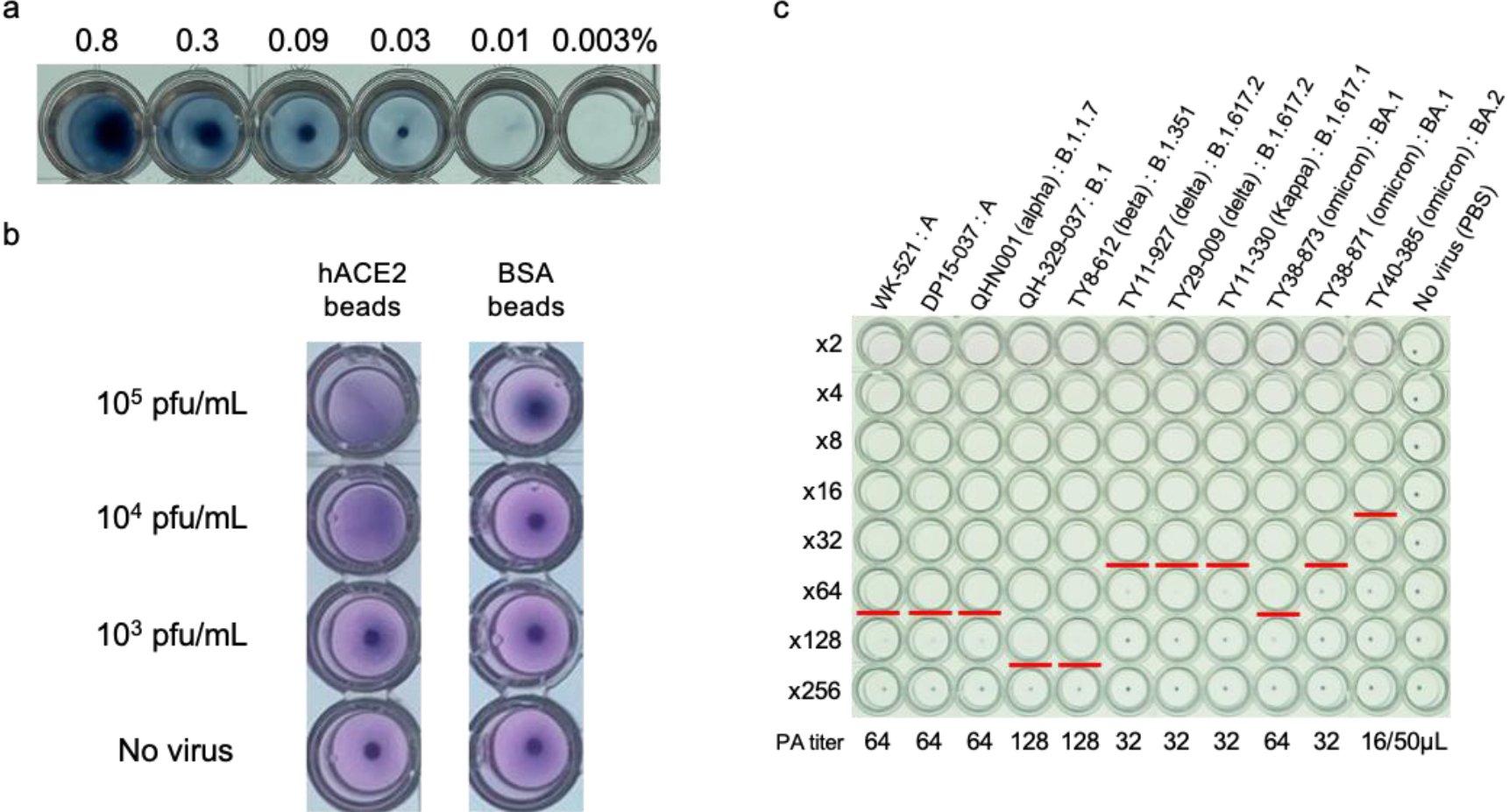
Establishment of the PA test. **a**, Optimization of hACE2-beads concentration. 2.5% hACE2-beads were diluted from 0.8~0.003% by PBS, and settled overnight at room temperature. **b**, Specificity of the PA test. SARS-CoV-2, QH-329-037 strain (50 μL), and 0.06% hACE2-beads/BSA-beads (50 μL) were mixed and settled overnight at room temperature. **c**, The PA test of the SARS-CoV-2 variants. A 2-fold dilution series of the variants (50 μL) was mixed with 0.01% hACE2-beads (50 μL), and then settled overnight at room temperature. The PA titer was defined as the highest dilution factor at which complete agglutination was observed (red line).

To confirm the specific binding of hACE2-beads to SARS-CoV-2, 50 μL of QH-329-037 isolate was mixed with 50 μL of 0.06% hACE2-beads (0.03% final concentration) and settled at room temperature. As a control, hACE2-unbound beads (BSA-beads) were prepared and tested in the same way as the hACE2-beads (Fig. 1b). A mixture of slight sedimentation and slight agglutination patterns was observed in the virus/hACE2-beads mixture after 6 h of settling, and the sedimentation and agglutination patterns could be clearly differentiated after overnight settling, whereas only a sedimentation pattern was observed in the virus/BSA-beads mixture. This suggests that SARS-CoV-2 can bind to hACE2-beads specifically and can be detected by observing the beads. To reduce the settling time required for the assay, another type of manufactured beads with a larger diameter (9.8 μm) were similarly prepared, and the PA test was carried out. However, although the settling time was reduced to <5 h, the 9.8 μm beads showed only a sedimentation pattern. Therefore, the 0.8 μm beads were used for the following assays. We also found that the sedimentation patterns were slightly harder to visualize on a V-bottom microtiter plate compared to a U-bottom plate, although the settling times on the two plates were almost the same. Therefore, a U-bottom plate was used for the subsequent assays. Thus, the tentative condition for the PA test was set as follows: mixing 50 μL of virus solution and 50 μL of 0.06% hACE2-beads (0.03% final concentration), followed by overnight settling at room temperature using a U-bottom plate. The results did not change when the beads were stored at 4°C for 1 month after preparation.

The PA test was performed on representative SARS-CoV-2 isolates under the tentative condition. Agglutination patterns were observed in all tested isolates, and the PA titer of the isolate was defined as the highest dilution factor at which complete agglutination was observed (Fig. S1a). However, the BA.2 omicron variant, TY40-385 isolate, showed the lowest PA titer (2PA unit/50 μL), which was inadequate to perform the following PAI test. To improve the PA titer, the concentration of hACE2-beads was re-examined. The final beads concentration was reduced from 0.03% to 0.005% and 0.0025%, and then the PA titers of three isolates (WK-521, TY11-927, and TY40-385) were compared among the three beads concentrations (Fig. S1b). All PA titers increased with declining hACE2-beads concentrations, and the final concentration of 0.005% was newly set as the standard condition based on the PA titer and visibility of sedimentation/agglutination patterns. Although sedimentation was not seen even at a final concentration of 0.01% in Fig. 1a, this was because the concentration of BSA was different between Fig. 1a and Fig. S1b. The retest with a 0.005% beads concentration resulted in an 8-fold increase in the PA titers of all isolates (Fig. 1c). The PA titers of isolates showed a strong positive correlation (r=0.85) with plaque-forming units measured by a plaque assay (Fig. S2), indicating that the PA titer reflects the amount of virus in the sample. However, the PA test showed different sensitivities among the variants and was the most sensitive to the omicron variants. This probably indicates that the omicron variants have a higher affinity for ACE2 than the previous variants^11^, and thus that they have a higher agglutination activity.

## Establishment of a particle agglutination inhibition (PAI) test

Next, the inhibitory effect of antibodies against SARS-CoV-2 on viral binding to hACE2-beads was evaluated based on the HI test for influenza virus6. We used antisera obtained from SARS-CoV-2-infected Syrian hamsters because these animals have been shown to be useful as a pathological model of SARS-CoV-2 infection, and the antibody titers against SARS-CoV-2 were increased in the infected hamsters^12^. Four hamsters per isolate were inoculated with the isolate (seven isolates in total), and the antisera were prepared. Elevation of antibody titer against the inoculated strain was confirmed by a cell-based micro-neutralizing test. As reported in the HI test for influenza virus6, nonspecific aggregation factor(s) that interfere with the inhibitory effect of antibodies were observed in some antisera at low dilution factors (Fig. 2a, sample 2). Therefore, prior to the use of such antisera, the factor(s) were removed by pre-adsorption treatment (see the Online Methods). The removal of the factor(s) from the antisera was confirmed by mixing the beads (Fig. 2a, sample 3).

**Figure 2.**
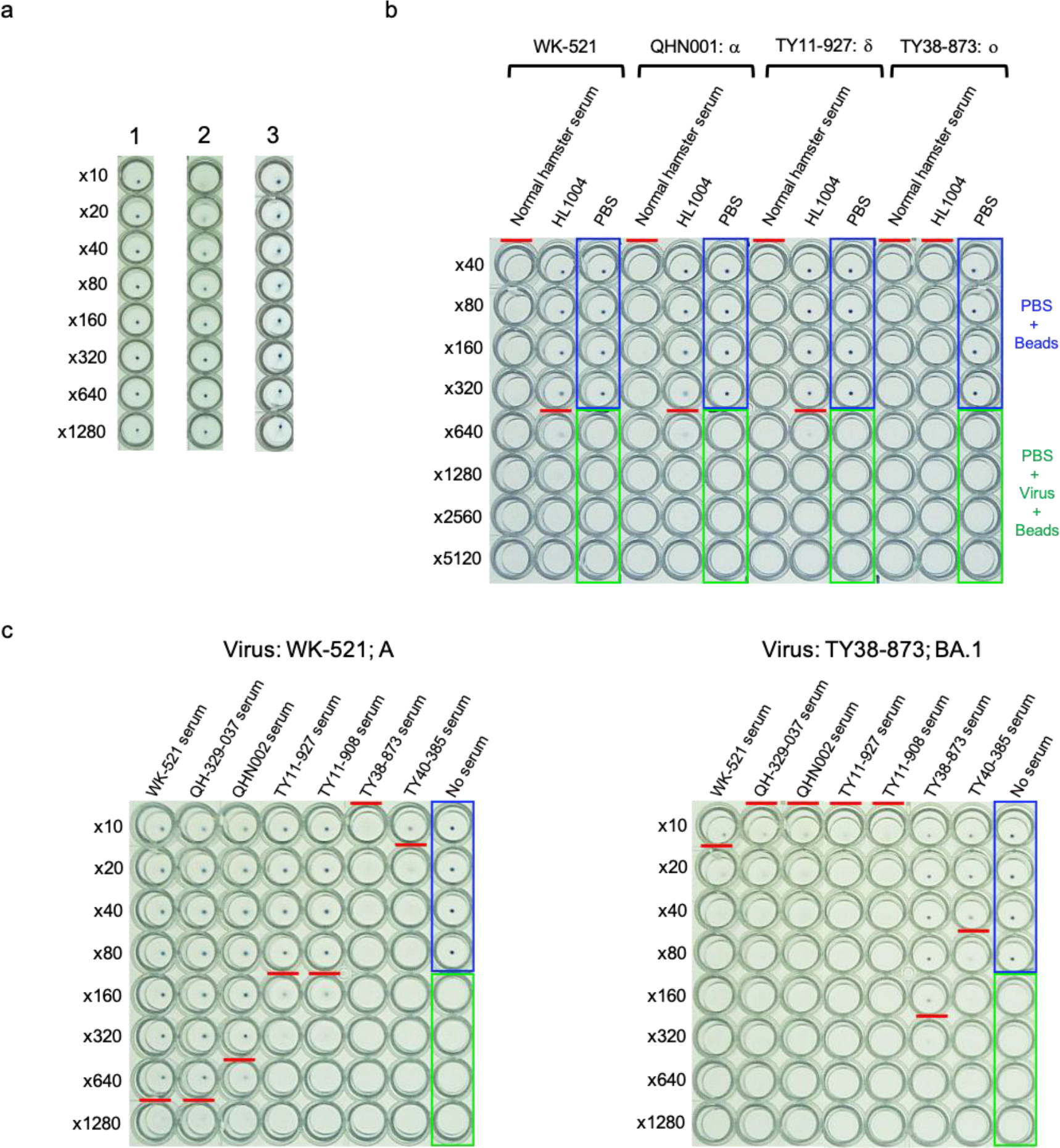
Establishment of the PAI test. **a**, The confirmation and removal of nonspecific agglutination factor(s). The serially diluted antisera (25 μL) were mixed with 0.01% hACE2-beads (25 μL), and then settled overnight at room temperature. 1: Antiserum without the factor(s); 2: antiserum with the factor(s); 3: antiserum after removal of the factor(s). **b**, Specificity of the PAI test. The serially diluted serum or antibody (25 μL) was mixed with the isolates (4PA/25 μL) and allowed to react for 60 min at room temperature. 0.01% hACE2-beads (50 μL) were added, and then settled overnight at room temperature. The wells framed in blue are the sedimentation controls with PBS instead of serum and virus. The wells framed in green are the agglutination controls with PBS instead of serum/antibody. The PAI titer is defined as the highest dilution factor at which complete agglutination inhibition was observed (red line). **c**, The PAI test of the SARS-CoV-2-infected antisera. Representative results are shown. The results for WK-521 (Pango Lineage: A) are shown at left, and the results for TY38-873 (BA.1) are shown at right.

The specificity of the binding inhibition was confirmed using a commercially available antibody, HL1004, which does not bind to the omicron variant BA.1 (GeneTex; catalog no. GTX635793: https://www.genetex.com/MarketingMaterial/Index/recombinant_antibodies_for_sars-cov-2_research; the web page does not list the reactivity for BA.1, but the same region as for the spike protein B.1.1.529 is used for the reactivity check). A normal (non-infected) Syrian hamster serum (Fujifilm Wako; catalog no. 569-76331) was also used a non-inhibitory control (Fig. 2b). Four PA units/25 μL of the isolates (WK-521 isolate; the ancestor, QHN001 isolate; the alpha variant, TY11-927 isolate; the delta variant and TY38-873 isolate; the omicron variant BA.1) was added to 25 μL of a 2-fold dilution series (from 40- to 5120-fold dilution) of the normal serum or the HL1004 antibody, and allowed to react for 60 min at room temperature. Then, 50 μL of 0.01% hACE2-beads was added (final concentration 0.005%) and settled overnight at room temperature. For the sedimentation or agglutination control, PBS was added in place of the antibody/serum or the isolates. PAI was not observed (all particles were agglutinated) with the normal hamster serum, whereas it was observed with the HL1004 antibody in the WK-521 isolate, the alpha and delta variants, and not in the omicron variant BA.1 as expected. The PAI titer was defined as the highest dilution factor of the serum/antibody for which complete agglutination inhibition was observed. The titers for the WK-521 isolate and the alpha, delta and omicron variants were 320, 320, 320 and <40, respectively. This suggests that hACE2-beads can be used for the PAI test under the following standard condition: mixing of 25 μL of serum and 25 μL of virus solution (4PA unit/25 μL) for 60 min, followed by mixing of 50 μL of 0.01% hACE2-beads, and overnight settling at room temperature.

The PAI test was then performed under the standard condition as defined above (Fig. 2c). Agglutination/agglutination-inhibition patterns were observed, reflecting the reactivity of the antiserum with the virus. The obtained PAI titers of each antiserum to the tested isolates are summarized in Table 1. The homologous PAI titer was defined as the titer against the same isolate/the same lineage of isolate used to prepare an antiserum as shown in Table 1 (highlighted in red). Anti-WK-521 serum showed a homologous PAI titer of 640, whereas the PAI titer was <10 against TY38-873 and *vice versa*: the homologous PAI titer and PAI titer were 160 and 10 against WK-521 in anti-TY38-873 serum. In the HI test for influenza viruses, antigenicity has generally been judged to be different when an HI titer is 8-fold or more different from a homologous HI titer. Based on this criterion, we defined an 8-fold or greater difference between the PAI titer and homologous PAI titer as an antigenic difference in our PAI test. Our results revealed that WK-521 (Lineage: A) was antigenically different from the beta (B.1.351) and omicron variants (BA.1 and BA.2). Similarly, QH-329-037 (B.1) and QHN002 (alpha, B.1.1.7) were antigenically different from the omicron variants (BA.1 and BA.2). The delta variants (TY11-927 and TY11-908, B.1.617.2) were antigenically different from the beta (B.1.351) and omicron variants (BA.1 and BA.2). The omicron variants were also antigenically different from the previous variants. This clearly showed that the antigenicity of the omicron and the previous variants/isolates are different. Antigenic differences between the D614G variant (QH-329-037 in this study) and the omicron variants (TY38-873, TY38-871 and TY40-385) have been reported in human^13^ or guinea pig^14^ convalescent and vaccinated sera by SARS-CoV-2 pseudo-virus neutralizing assay. Furthermore, the results of the PAI test also suggested antigenic differences between the WK-521 isolate and the beta variant (TY8-612), and between the beta and delta variants (TY11-927 and TY11-908). The antigenic difference of the beta variant has been reported by a structure-function analysis of the monoclonal antibodies from beta-variant-infected individuals15. Collectively, our findings indicate that the PAI test is a useful method for the antigenic analysis of SARS-CoV-2, and is easier than the previously available methods.

**Table 1.** The PAI titers of SARS-CoV-2-infected hamster antisera.

|  | WK-521<br>A<br>Serum | QH-329-037<br>B.1<br>Serum | QHN002<br>Alpha : B.1.1.7<br>Serum | TY11-927<br>Delta : B.1.617.2<br>Serum | TY11-908<br>Delta : B.1.617.2<br>Serum | TY38-873<br>Omicron : BA.1<br>Serum | TY40-385<br>Omicron : BA.2<br>Serum |
| --- | --- | --- | --- | --- | --- | --- | --- |
| WK-521<br>A | 640 | 640 | 320 | 80 | 80 | < 10 | 10 |
| DP15-037<br>A | 640 | 640 | 320 | 80 | 80 | <10 | 10 |
| QH-329-037<br>B.1 | 640 | 640 | 640 | 160 | 160 | < 10 | 10 |
| QHN001<br>Alpha : B.1.1.7 | 640 | 640 | 640 | 80 | 80 | <10 | 10 |
| TY8-612<br>Beta : B.1.351 | 80 | 320 | 320 | < 10 | 20 | < 10 | 10 |
| TY11-927<br>Delta : B.1.617.2 | 320 | 320 | 160 | 160 | 160 | < 10 | < 10 |
| TY29-009<br>Delta : B.1.617.2 | 160 | 160 | 160 | 80 | 80 | < 10 | < 10 |
| TY11-330<br>Kappa : B.1.617.1 | 320 | 320 | 160 | 80 | 40 | < 10 | < 10 |
| TY38-873<br>Omicron : BA.1 | 10 | < 10 | < 10 | < 10 | < 10 | 160 | 40 |
| TY38-871<br>Omicron : BA.1 | < 10 | < 10 | < 10 | < 10 | < 10 | 80 | 40 |
| TY40-385<br>Omicron : BA.2 | 20 | 40 | 40 | < 10 | < 10 | 10 | 160 |

The homologous PAI titers are shown in red.

## Discussion

In this study, we established the PA/PAI test using hACE2-coated beads. The PA titers were correlated with the amount of SARS-CoV-2. The PAI titers clearly showed the antigenic differences between the omicron and the previous variants. Furthermore, the titers also suggested antigenic differences of the beta variant from the WK-521 (original Wuhan strain) isolate and the delta variant. These antigenic differences were supported by the results shown in previous reports^13–15^.

The PA/PAI methods allow titration of SARS-CoV-2 and antibodies to the virus *via* a much simpler process than traditional cell-based assays, such as plaque assay or neutralization assay. This is beneficial for the continuous monitoring of antigenicity of field isolates in SARS-CoV-2 surveillance. In addition, the PAI assay could be used for the evaluation of antibody titers in individuals.

Current vaccines of SARS-CoV-2 using the spike protein or its gene are based on the Wuhan-Hu-1 strain. However, this study showed that the omicron variants possess different antigenic properties. Continuous monitoring of the viral antigenicity of field isolates and the holding status of protective antibodies in certain populations will be a crucial component of SARS-CoV-2 surveillance. A review of vaccine strains will also be necessary. The PA/PAI method for SARS-CoV-2 established in this study should be a useful tool to obtain informative data for such discussions.

As shown above, the PA/PAI assay is an easier method, but its cost might be a concern. This assay can be performed for ~US $30 per 96-well plate. This study used commercially available hACE2 to save the time to prepare the protein, which was the most expensive (~US$800/100 μg). If hACE2 can be prepared in-house, the cost can be further reduced to ~US $1 per 96-well plate. Finally, we note that a PA/PAI test using inactivated SARS-CoV-2 was not performed in this study. If the PA/PAI test were to be validated with inactivated SARS-CoV-2, the assay could be used without any restriction due to the biosafety level.

## Author contributions

H.H., S.W. and K.N. conceived the project. J.K., S.M. and K.N. designed the study. J.K. performed the beads preparation, PA/PAI tests and SARS-CoV-2 culture. S.M. performed the SARS-CoV-2 culture, the first PA test and the plaque assay. Y.S., T.A. M.S., H.A. and S.W. prepared the hamster sera. J.K., S.M., S.W and K.N. wrote the manuscript. All the authors commented on the manuscript.

## Acknowledgments

We are grateful to Hiyori Okura for excellent technical assistance. This work was partially supported by a Grant-in-Aid for Emerging and Reemerging Infectious Diseases from the Ministry of Health, Labor, and Welfare of Japan (no. 20HA2007), AMED Grants (no. 21nf0101626j0102) to H.H., and AMED Grants (no. 21fk0108104k0003), a JSPS grant (no. 20K07519) and a Uehara Memorial Foundation grant (no. 202120234) to S.M.

## Online Methods

### SARS-CoV-2 culture

80~95% confluent Vero E6 cells expressing transmembrane protease serine 2 (TMPRSS2)^16^ were infected with SARS-CoV-2 strains using DMEM supplemented with 5% fetal bovine serum and penicillin/streptomycin, and then cultured at 37oC under 5% CO_2_. WK-521 (Pango lineage: A), DP15-037 (Pango lineage: A), QH-329-037 (Pango lineage: B.1), QHN001 (alpha variant, Pango lineage: B.1.1.7), TY8-612 (beta variant, Pango lineage: B.1.351), TY11-927 (delta variant, Pango lineage: B.1.617.2), TY29-009 (delta variant, Pango lineage: B.1.617.2) and TY11-330 (kappa variant, Pango lineage: B.1.617.1) isolates were harvested 24 h after infection by centrifugation at 1000 rpm for 5 min at room temperature. TY38-873 (omicron variant, Pango lineage: BA.1), TY38-871 (omicron variant, Pango lineage: BA.1) and TY40-385 (omicron variant, Pango lineage: BA.2) isolates were harvested 48 h after infection by centrifugation. The harvested viruses were stored at −80°C until use.

### Beads (particle) preparation

Deep blue-dyed latex beads, 0.8 μm in average diameter, were purchased from Sigma-Aldrich (catalog no. L1398). 0.5 mL of 2.5% (w/v) beads was centrifuged at 2400g for 10 min at room temperature, and the beads were recovered as precipitates. The beads were washed twice with 1 mL PBS (Takara; catalog no. T900) using centrifugation. The washed beads were centrifuged and resuspended in 0.25 mL of 25 mM MES-NaOH, pH 6.0. The resuspended beads were centrifuged and mixed with 2.5 mL of 25 mM MES-NaOH, pH 6.0 containing 100 μg hACE2 (GeneTex; catalog no. GTX01550-pro) for 24 h at 4°C using a rotator. The hACE2-bound beads were centrifuged at 2400g for 10 min at 4°C, and washed twice with 1 mL PBS. The OD280 of the hACE2 solution was measured before and after mixing of hACE2 solution and the beads. The beads were blocked with 0.75 mL of PBS with 3% bovine serum albumin (BSA, Sigma-Aldrich; catalog no. A9418) for 30 min at room temperature. The blocked beads were stored in 0.5 mL of PBS with 1% BSA (final 2.5% (w/v) beads concentration) at 4°C until use.

### PA test

2.5% (w/v) hACE2-beads were diluted to 0.01% (w/v) with PBS supplemented with 1% BSA. 50 μL aliquots of a 2-fold dilution series of SARS-CoV-2 variants were prepared in a 96-well plate with PBS. 50 μL of 0.01% (w/v) hACE2-beads was added to each well. After overnight settling at room temperature, sedimentation/agglutination patterns were observed. A mixture of 50 μL of PBS and 50 μL of the beads was used as the sedimentation (no-agglutination) control.

### Titration of SARS-CoV-2

The titers of SARS-CoV-2 used in this study were determined either by plaque assay or by 50% tissue culture infectious dose (TCID_50_). For the plaque assay, monolayers of VeroE6/TMPRSS2 cells grown in a 96-well plate were infected with serially diluted culture supernatants of SARS-CoV-2, cultured in high-glucose Dulbecco's modified Eagle's medium (DMEM; Sigma-Aldrich) containing 2.5% carboxymethyl cellulose at 37oC under 5% CO_2_ for 3 days, and then fixed with 4% paraformaldehyde and stained with crystal violet. Emergent plaques were counted using an optical microscope. For TCID_50_, 10-fold serially diluted viruses were mixed with VeroE6/TMPRSS2 cells (2-3 x 104 cells/well) in a 96-well plate and incubated at 37°C under 5% CO_2_ for 5 days. Five days later, the cytopathic effect in each well was checked and the TCID_50_ was determined by the Kärber method^17^.

### Preparation of hamster antisera and micro-neutralizing test

The five-week-old female Syrian hamsters were intranasally inoculated with 50 μL of 10^3^ TCID_50_ of each SARS-CoV-2 strain (WK-521, QH-329-037, QHN002, TY11-927, TY11-908, TY38-873 and TY40-385). The whole blood was collected by cardiac puncture under deep terminal anesthesia 14-16 days after infection and sera were prepared by centrifugation. The sera were inactivated at 56oC for 30 min to use for the microneutralization and the PAI test.

Two-fold serial dilutions of sera were mixed with 10^2^ TCID_50_ of the SARS-CoV-2 strain and pre-incubated in 96-well plates at 37°C for 60 min. After pre-incubation, VeroE6/TMPRSS2 cells were added to the virus-serum mixture and incubated at 37°C under 5% CO_2_ for 5 days. Five days later, the cytopathic effect in each well was checked and the microneutralization titers of sera were determined as the reciprocal of the highest dilution that did not display the cytopathic effect.

### PAI test

Prior to performing the PAI test, the nonspecific agglutination factor(s) in antisera was removed if antiserum samples showed nonspecific agglutination as follows. 500 μL of 2.5% hACE2-beads was centrifuged and the supernatant was removed. 1 mL of 10-fold diluted antiserum with PBS was added to the precipitated beads, and then mixed by a rotator for 60 min at room temperature. The treated antiserum was centrifuged at 2400g for 10 min at 4°C and the collected supernatant was used for the following PAI test.

25 μL aliquots of 2-fold dilution series (from 10- to 1280-fold) of each antiserum were prepared in 96-well plates with PBS. 4PA units/25 μL of SARS-CoV-2 isolates were added to the diluted antisera and incubated for 60 min at room temperature. The accuracy of the PA titer of the added viruses was confirmed by another PA test (back titration). 50 μL of 0.01% (w/v) hACE2-beads were added to each well. After overnight settling at room temperature, the agglutination/agglutination-inhibition patterns were observed. For the sedimentation or agglutination control wells, 25 μL PBS was used in place of antisera or virus, respectively.

