## Supplemental Figures for "Use of the particle agglutination/particle agglutination-inhibition test for antigenic analysis of SARS-CoV-2"

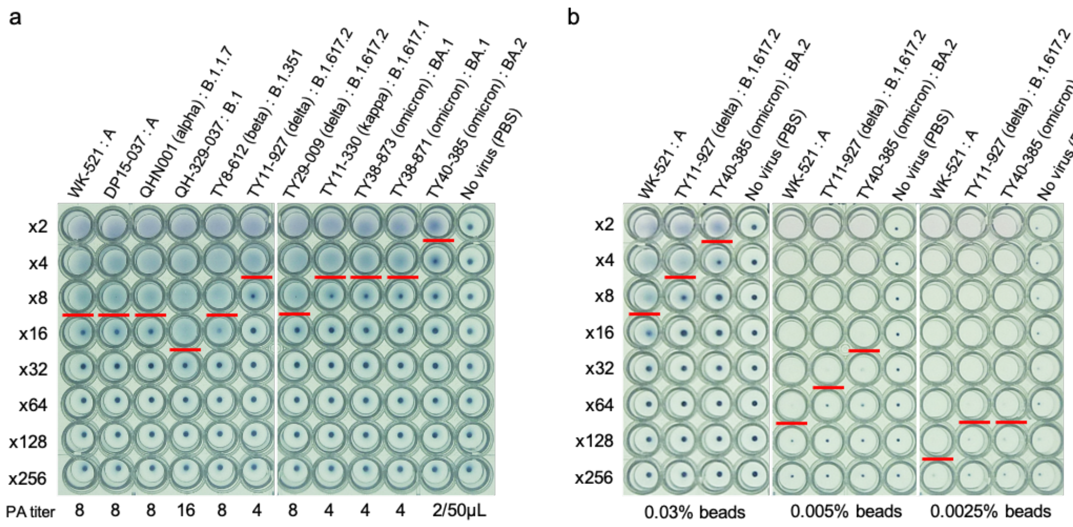


Figure S1. Optimization of the PA test.

**a**, The result using 0.03% hACE2-beads. A 2-fold dilution series of the viruses (50 μL) was mixed with 0.06% hACE2-beads (50 μL), and then settled overnight at room temperature. **b**, Optimization of the hACE2-beads concentration. The final beads concentration was varied from 0.03~0.0025%. The PA titers are shown with red lines.

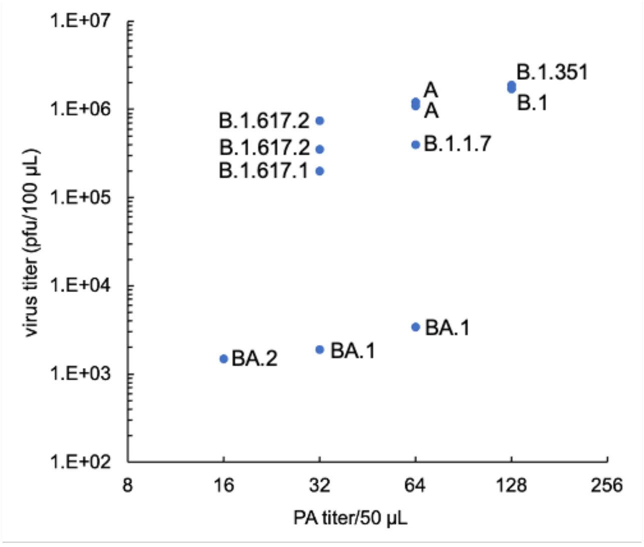


Figure S2. The correlation between the PA titer and the plaque-forming units.
